# Right hemisphere superiority for executive control of attention

**DOI:** 10.1101/432732

**Authors:** Alfredo Spagna, Tae Hyeong Kim, Tingting Wu, Jin Fan

**Affiliations:** Department of Psychology, Columbia University in the city of New York, NY, USA; institute du Cerveau et de la Moelle epiniere, ICM, Inserm U 1127, CNRS UMR 7225, Sorbonne Universite, Paris, France; Department of Psychology, Queens College, the City University of New York, Flushing, NY, USA; Departments of Psychiatry and Neuroscience, Icahn School of Medicine at Mount Sinai, New York, NY, USA

## Abstract

Over forty years have passed since the first evidence showing the unbalanced attentional allocation of humans across the two visual fields, and since then, a wealth of behavioral, neurophysiological, and clinical data increasingly showed a right hemisphere dominance for orienting of attention. However, inconsistent evidence exists regarding the right-hemisphere dominance for executive control of attention, possibly due to a lack of consideration of its dynamics with the alerting and orienting functions. In this study, we used a version of the Attentional Network Test with lateralized presentation of the stimuli to the left visual field (processed by the right hemisphere, RH) and right visual field (processed by the left hemisphere, LH) to examine visual field differences in executive control of attention under conditions of alerting or orienting. Analyses of behavioral performance (reaction time and error rate) showed a more efficient executive control (reduced conflict effect) in the RH compared to the LH for the reaction time, under conditions of increased alerting and of informative spatial orienting. These results demonstrate the right-hemisphere superiority for executive control, and that this effect depends on the activation of the alerting and orienting functions.

## Introduction

Executive control of attention plays a central role in human cognition as the mechanism that allows the selection and prioritization of the processing of goal-relevant information to reach consciousness, and is achieved via the interplay with the other two components of attention, the alerting function for the preparation and maintenance of a state of readiness, and the orienting function for the direction of attention towards relevant features of a stimulus (Fan et al., 2009; Petersen & Posner, 2012; Spagna, Mackie, & Fan, 2015). Evidence in favor of a right hemisphere dominance for orienting of attention came initially from studies of patients with unilateral hemispatial neglect (Bartolomeo, 2007; Bartolomeo & Chokron, 2002; Chica et al., 2012; Heilman & Van Den Abell, 1980; Kinsbourne, 1987; Lunven & Bartolomeo, 2016; Mesulam, 1999; Posner, Cohen, & Rafal, 1982; Posner, Walker, Friedrich, & Rafal, 1984), and were later supported by most of the behavioral patterns showing better performance in orienting to stimuli presented in the left visual field (processed by the right hemisphere, RH) compared to stimuli presented in the right visual field (processed by the left hemisphere, LH) in healthy individuals (De Schotten et al., 2011; Kincade, Abrams, Astafiev, Shulman, & Corbetta, 2005; Marotta, Lupianez, & Casagrande, 2012; Shulman et al., 2009; Smigasiewicz, Westphal, & Verleger, 2017; Vossel, Weidner, Driver, Friston, & Fink, 2012; Zago et al., 2017; Zuanazzi & Cattaneo, 2017). Because the purpose of orienting is to ultimately facilitate the processing of information and conflict resolution (Callejas, Lupianez, Funes, & Tudela, 2005; Callejas, Lupianez, & Tudela, 2004; Fan et al., 2009) by focusing more on the task-relevant feature or location of the imperative stimuli and therefore achieve cognitive control (Mackie, Van Dam, & Fan, 2013), theoretically there should be a RH superiority for the executive control of attention. However, there is inconsistent evidence regarding the right-lateralization of the other specific functions of attention, especially for the executive control (Asanowicz, Marzecova, Jaskowski, & Wolski, 2012; Greene et al., 2008; Konrad et al., 2005), with some studies showing better conflict resolution in the RH compared to the LH (Asanowicz et al., 2012; Garavan, Ross, & Stein, 1999; Mecklinger, von Cramon, Springer, & Matthes-von Cramon, 1999; Spielberg et al., 2011), but not in others (Greene et al., 2008; Konrad et al., 2005; Spagna, Martella, Fuentes, Marotta, & Casagrande, 2016; Wu, Weissman, Roberts, & Woldorff, 2007). Therefore, whether a RH advantage exists for executive control remains unclear, and determining the laterality of this function will further support the notion of RH dominance for attention (Heilman & Van Den Abell, 1980; Kinsbourne, 1987; Mesulam, 1981; Weintraub & Mesulam, 1987).

The pivotal role of the RH in the allocation of attentional resources, especially in the visuospatial environment, was initially (and consistently throughout the past 40 years) shown by studies in patients with unilateral hemispatial neglect, a neurological disorder that often follows a lesion to the right parietal cortex and / or underlying white matter which results in patients failing to direct their attention to the contra-lesional (i.e, left) side of the space (Bartolomeo & Chokron, 2002; Chica, Thiebaut de Schotten, Bartolomeo, & Paz-Alonso, 2018; Corbetta & Shulman, 2011; Heilman & Van Den Abell, 1980; Lunven & Bartolomeo, 2016; Mesulam, 1999; Posner et al., 1984; Rafal, 1994; Rastelli et al., 2013; Toba et al., 2018). This deficit in the ability to select information (in other terms “orient”) for goal-directed behavior following RH damage has been studied in a great deal of literature regarding the interaction between hemisphere and this function of attention (e.g., Benwell, Thut, Grant, & Harvey, 2014; Chica et al., 2012; Foxe, McCourt, & Javitt, 2003; Longo, Trippier, Vagnoni, & Lourenco, 2015; Marotta et al., 2012; Shulman et al., 2010; Spagna et al., 2016). Based on this evidence, the *hemispatial theory* (Heilman & Van Den Abell, 1980) and the *interhemispheric competition account* (Kinsbourne, 1987) are two models proposing alternative mechanisms underlying the RH advantage of attention: while the former stated that the RH advantage is rather a “disadvantage” of the LH that is able to orient only towards the contralateral visual field (as opposed to the RH that can direct attention to both visual fields, hence the dominance), the latter proposed that each hemisphere has its own “contralateral vector of attention”, and that a lesion to one of the two hemispheres disrupts the normal balance and favors the orienting towards the ipsilesional visual field. Perhaps due to the involvement of both hemispheres in orienting, as shown in a wealth of neurophysiological studies (Corbetta, Patel, & Shulman, 2008; Corbetta & Shulman, 2002; Fan, McCandliss, Fossella, Flombaum, & Posner, 2005; Xuan et al., 2016), behavioral studies have revealed inconsistent results concerning this function of attention, with some studies revealing a RH dominance (exp. 1 of Asanowicz et al., 2012; Evert, McGlinchey-Berroth, Verfaellie, & Milberg, 2003; Greene et al., 2008; Kalamala, Drozdzowicz, Szewczyk, Marzecová, & Wodniecka, 2018; Poynter, Ingram, & Minor, 2010; Śmigasiewicz, Asanowicz, Westphal, & Verleger, 2014), while other studies failed to find a hemispheric asymmetry (exp. 2 of Greene et al., 2008; Spagna et al., 2016; Tao, Marzecová, Taft, Asanowicz, & Wodniecka, 2011). Controversies also exist regarding the RH superiority of the other two components, the alerting and executive control function. Behavioral studies have not found significant visual field difference for the alerting network in healthy individuals (Asanowicz et al., 2012; Greene et al., 2008; Marzecová, Asanowicz, KrivÁ, & Wodniecka, 2012; Spagna et al., 2016), possibly due to the simplicity of the testing procedure used, as stronger attentional asymmetries have been shown to come from more difficult tasks (Jonides, 1979; Verleger et al., 2009; Welcome & Chiarello, 2008). Nonetheless, a greater deficit in alerting has been found after RH damage to regions of the parietal lobe partially overlapping with those related to the hemispatial neglect discussed above (Fernandez-Duque & Posner, 2001; Petersen & Posner, 2012; see also Posner, 2008). For the executive control, behavioral evidence of an hemispherical asymmetry mostly derives from a lateralized version of the Stroop task (Stroop, 1935), a task that heavily relies on the well-known left-lateralized language function, and a RH advantage for conflict processing has been found in some studies (Asanowicz et al., 2012; Kalamala et al., 2018; Marzecová et al., 2012; Poynter et al., 2010; Weekes & Zaidel, 1996) but not in others (Belanger & Cimino, 2002; Greene et al., 2008; Konrad et al., 2005; Spagna et al., 2016). Overall, whether a RH dominance of attention exists across all three functions remains unclear, along with the mechanisms responsible for such unilateral advantage of attention.

A potential reason for the above-mentioned inconsistent results, especially regarding the executive control function, may reside in the neglected consideration of the interplay among the components (Badre, 2011) that leads to the unitary construct of attention. Although initially conceived as three independent components, both in terms of function (Fan, McCandliss, Sommer, Raz, & Posner, 2002) and associated brain structure (Fan et al., 2005), the interplay between the alerting, orienting, and executive control was soon after identified as a key component in the prioritization of mental computations (Callejas et al., 2005; Callejas et al., 2004; Chica et al., 2012; Fan et al., 2009; Liu, Bengson, Huang, Mangun, & Ding, 2016; Macaluso, 2010; Martella, Casagrande, & Lupianez, 2011; Spagna, Dong, et al., 2015; Spagna, Mackie, et al., 2015; Spagna et al., 2014). For example, efficient dynamics among these functions may ameliorate the inattentional symptoms associated with hemispatial neglect (Chica et al., 2012), which may open critical rehabilitation perspectives (Manly, Hawkins, Evans, Woldt, & Robertson, 2002; Robertson, Manly, Andrade, Baddeley, & Yiend, 1997). Evidence of modulatory effects involving the alerting function were first observed in studies showing the beneficial effect of an alerting cue on the orienting function (Fernandez-Duque & Posner, 1997; Fuentes & Campoy, 2008; Q. Li, Liu, Huang, & Huang, 2018; Mullane, Lawrence, Corkum, Klein, & McLaughlin, 2016; Wiegand, Petersen, Bundesen, & Habekost, 2017), and the detrimental effect of an alerting cue (whether visual or auditory) in conflict resolution (Asanowicz & Marzecová, 2017; Callejas et al., 2005; Callejas et al., 2004; Zani & Proverbio, 2017). The synergistic cooperation between the endogenous (i.e., voluntary) orienting and executive control function has also been shown in the form of a more precise selection of target information reducing the distracting effect of irrelevant information (Callejas et al., 2005; Callejas et al., 2004; Fan et al., 2009; Fuentes & Campoy, 2008; Spagna, Mackie, et al., 2015), while exogenous (i.e., automatic) orienting has been shown to impair conflict resolution (Trautwein, Singer, & Kanske, 2016). Executive control has been proposed to be located at the top of a hierarchical structure and acting irrespectively of sensory modalities (i.e., supramodal) (Donohue, Liotti, Perez, & Woldorff, 2012; Moris Fernandez, Macaluso, & Soto-Faraco, 2017; Roberts & Hall, 2008; Spagna et al., 2017; Spagna, Mackie, et al., 2015), with the alerting and orienting functions being located at a lower level and tied more to modality-specific mechanisms (Bushara et al., 1999; Langner et al., 2011; Salmi, Rinne, Degerman, Salonen, & Alho, 2007; Thiel & Fink, 2007; Ward, 1994; Yang & Mayer, 2013). Therefore, investigating the hemispherical asymmetries of the executive control under different states of alerting and orienting may provide useful insights on the dynamics from which attention emerges as a unitary cognitive function.

In this study we employed the visual field methodology (Bourne, 2006) to examine the efficiency and interactions of the attentional networks separately in the RH and LH. The hemispherical asymmetry of the executive control was investigated under different alerting and orienting conditions by using a lateralized version of the attentional network test (LANT-R). In this task, the hemispherical difference for executive control of attention was examined by presenting the target in the left or right visual field, under the conditions of congruent or incongruent flankers to generate the conflict effect, preceded by bilateral or unilateral visual cues to trigger the alerting and the orienting functions, respectively. We predicted a RH superiority in the executive control function, as shown by a reduced conflict effect, and that such superiority should be facilitated by the activation of alerting and orienting functions.

## Method and Materials

### Participants

Fifty students taking the Introduction to Psychology course at Queens College, the City University of New York (CUNY) participated in this study. Data from two participants were not included in the analyses due to low accuracy (approaching chance level) and an overall response time (RT) greater than two standard deviations (SD) from the group mean. The remaining participants (n = 48) consisted of 40 females and 8 males, with an average age of 20.8 years old (SD = 3.22), ranging from 18 to 25. All but one participant were right-handed, and all participants had normal or corrected-to-normal vision. Written informed consent approved by the Institutional Review Board of Queens College, CUNY was obtained from all participants prior to participation.

### The Lateralized Attention Network Test - Revised (LANT-R)

The LANT-R is a modified version of the revised attention network test (Fan et al., 2009) to measure the hemispherical differences in the efficiency of the attentional functions (alerting, orienting, executive control) and their interactions. **Figure 1** illustrates the sequence of events of the LANT-R. On the screen with a gray background, there were two vertically aligned rectangular boxes with a black outline located to the left and right of a central fixation cross. For each trial, five arrows appeared in either the left or right box, with the center arrow (the target) pointing either up or down, and the other arrows located above or below the target pointing either toward the same direction (flanker congruent condition) or toward the opposite direction (flanker incongruent condition). Participants were required to indicate the direction of the target by pressing the corresponding button on the mouse. The target was cued under one of three cueing conditions: double cue (i.e., the outline of both boxes changing from black to white), spatial cue (the outline of one of the boxes changing from black to white), or no cue (no change in the outline of any of the boxes). The double cue was used as an alerting stimulus by providing temporal information about the impending target, regardless of location. The spatial cue was designed to validly or invalidly orient the participant’s attention to either the left or the right side, thus providing both temporal and spatial information about the impending target. In each trial, participants were required to indicate the direction of a central arrow surrounded both above and below by two flanker arrows, either pointing in the same (congruent) or in the opposite (incongruent) direction as the target.

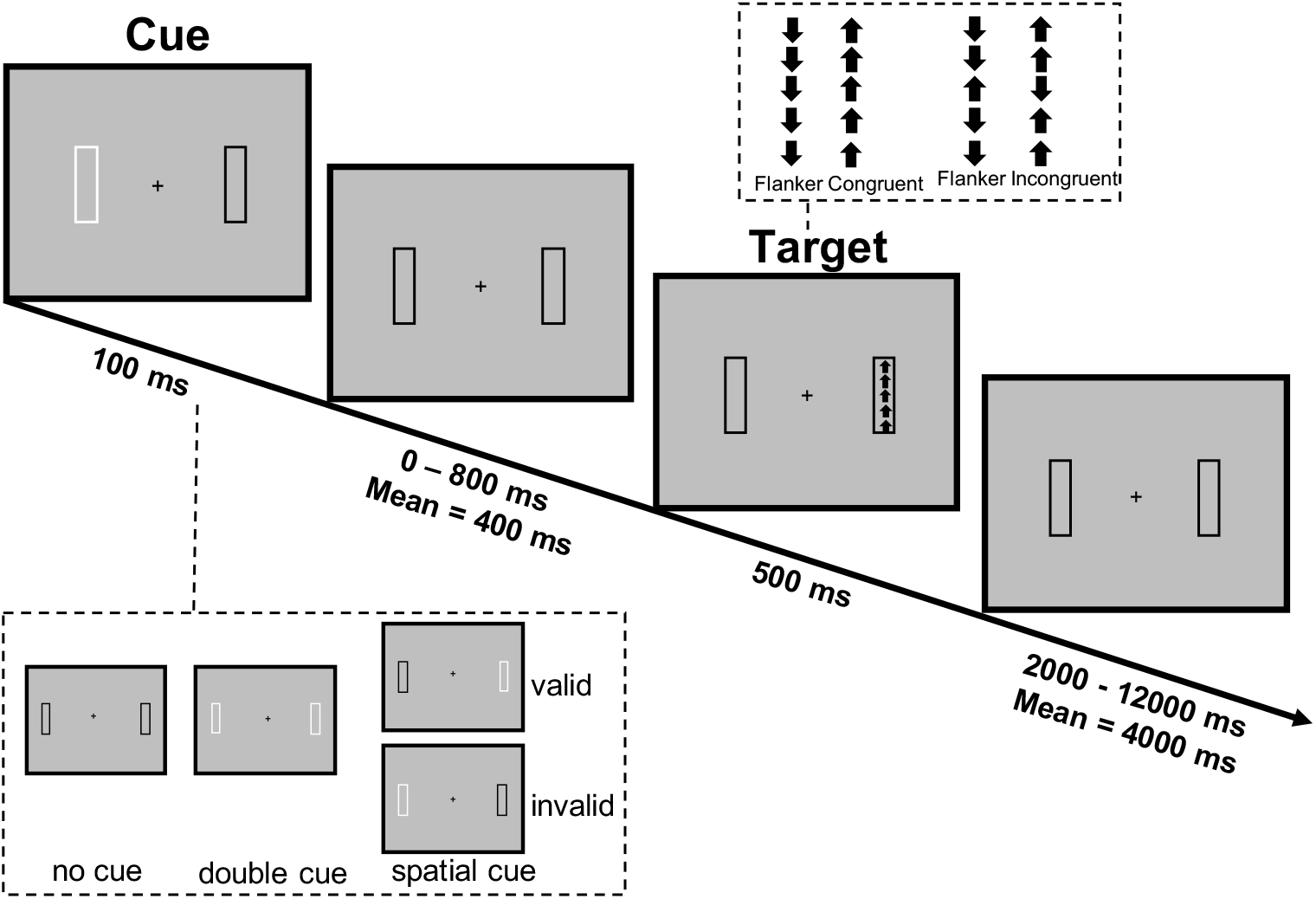

The size of the central fixation cross, located in the middle of the screen and visible throughout the entire task, was 1° of visual angle. The five black arrows that appeared on either side were .56° each. The arrows were separated by .06° of space, and the stimuli (target arrow and the flankers) subtended a total of 3.27°. Each box was located 5° from the central fixation cross. Preceding the target by three cue-to-target intervals (0 ms, 400 ms, 800 ms) equally likely to be presented, the change in the outline of the boxes from black to white lasted 100 ms, then the target and flankers were presented for 500 ms. Following the arrow presentation, there was a variable inter-trial interval ranging from 2000 to 12000 ms, with a mean interval of 4000 ms. Each trial had a mean duration of 5000 ms. The test consisted of 4 blocks, with each block containing 72 trials. On half of the trials (i.e., 144), the participant was shown a valid spatial cue, while the remaining trials were equally divided in 48 trials of a double cue condition, 48 trials of an invalid spatial cue condition, and 48 trials of a no cue condition. There was equal number of trials in the congruent and incongruent condition. In total, each block took 420 seconds to be completed, and the entire experiment took around 30 minutes to be completed. Prior to the beginning of the experimental session, the participant completed a short practice session consisting of 32 trials. During the practice session, participants were provided with feedback on accuracy and response time on each trial.

The practice session and the experimental session provided instructions in written form on the computer screen. It was emphasized that the participants fixate on the central fixation cross for the duration of each trial and press the mouse button that corresponded to the direction of the target arrow as quickly and accurately as possible. The mouse was tilted 90 degrees to the left, so that the right mouse button indicated “up”, and the left mouse button indicated “down”. The mouse was also aligned to the middle of the computer screen and participant’s midline of the body. To diminish the effects of hemispheric bias through motor control, the participant used both hands on the mouse to respond (Gable, Poole, & Cook, 2013): the right index finger was used to press the up-button while the left index finger was used to press the down-button. The task was displayed on a 17-inch LCD monitor, and the LANT-R was programmed in E-Prime (Psychology Software Tools, Pittsburgh, PA). Each participant completed the testing on a computer located in a silent and well-illuminated room.

### Operational definition of the attentional effects and interactions

A 4 × 2 × 2 factorial design was used in this experiment, with *cue condition* (no cue, double cue, valid cue, invalid cue), *conflict condition* (congruent, incongruent), and *visual field* (left, right) as within-subject factors. The same operational definitions for the attentional functions presented originally in (Fan et al., 2009) were used to estimate the attentional effects and are shown in **Table 1**. The *alerting effect* represents the performance benefit produced by the increased arousal compared to the no cue condition. The *orienting effect* is equivalent to *moving + engaging* operation defined in (Fan et al., 2009) and represents the performance benefit produced by a valid spatial information compared to the temporal information provided by an alerting cue. The *disengaging effect* represents the cost of disengaging from an invalid spatial cue. Combining the *disengaging* operation and the *moving + engaging* operation leads to the *validity effect*, which represents the extent to which a valid spatial cue condition benefits the participant’s performance compared to the cost in performance due to the invalid spatial cue condition. The *conflict effect* represents the cost in solving the conflict caused by the incongruent flankers, and a larger conflict effect corresponds to a less efficient executive control function (Fan et al., 2002). The interaction effects between these attentional functions were calculated by comparing the conflict effects under different cue conditions: *alerting by conflict interaction effect*, with a negative value indicating a negative impact of alerting on conflict processing. *Orienting by conflict interaction effect*, with a positive value indicating a more efficient conflict processing because of orienting. *Disengaging by conflict interaction effect*, with a positive value indicating a less efficient conflict processing because of invalid orienting compared to an alerting cue. *Validity by conflict interaction effect*, with a positive value indicating a less efficient conflict processing because of invalid orienting compared to a valid orienting cue. All the above-defined effects and interactions were computed separately for the RH and LH, according to the location where target and flankers were presented.

**Table 1.**
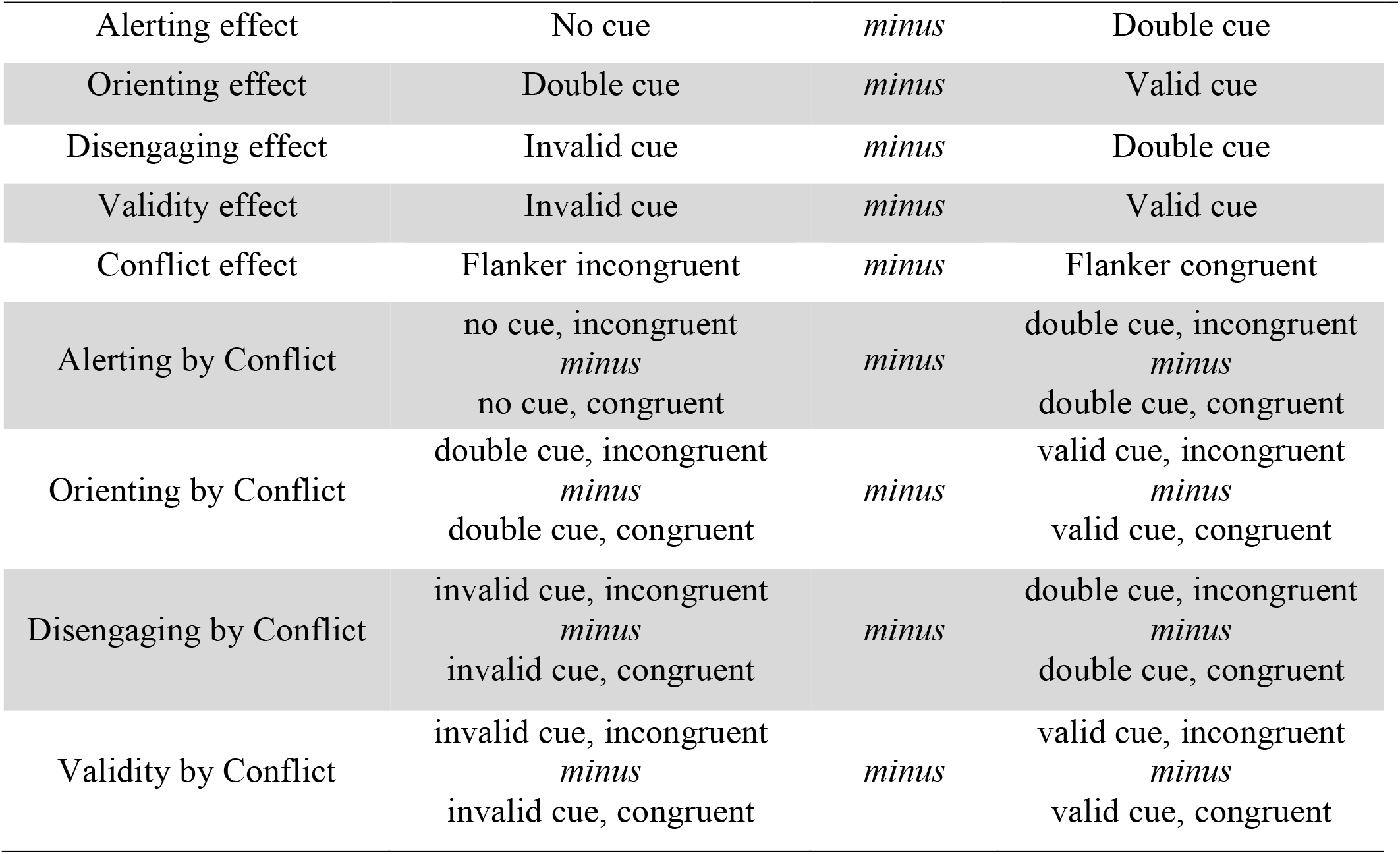
Operational definition of the attentional effects and interactions

### Data Analyses

Mean RT and error rate were calculated for each condition, with error trials (incorrect or missing responses) (less than 2% per subject) and RT outliers (above two standard deviations) (less than 2.23% per subject) being excluded from analyses on RT. Hemispherical differences in attentional effects, overall RT, and error rate, were tested using paired-sample *t* tests (one-tailed, hypothesizing a RH advantage). Effect sizes are reported as Cohen’s *d.* Further, a 4 × 2 × 2 ANOVA was conducted to test for the presence of *omnibus* effects and interactions among cue conditions, congruent conditions, and the visual fields. Planned comparisons were then conducted to analyze significant interactions between the conflict effect and the alerting, orienting, disengaging, and validity effects were tested using 2 × 2 × 2 ANOVAs, with an alpha-level set to .05. Specifically, a 2 (*Hemisphere:* RH, LH) × 2 (*Alerting:* no cue, double cue) × 2 (*Conflict:* congruent, incongruent) was conducted to identify differences in the alerting × conflict interaction in the two hemispheres. A 2 (*Hemisphere:* RH, LH) × 2 (*Orienting:* double cue, valid cue) × 2 (*Conflict:* congruent, incongruent) was conducted to examine the orienting × conflict interaction separately in each hemisphere. A 2 (*Hemisphere:* RH, LH) × 2 (*Disengaging:* invalid cue, double cue) × 2 (*Conflict:* congruent, incongruent) was conducted to examine the disengaging × conflict interaction separately in each hemisphere. A 2 (*Hemisphere:* RH, LH) × 2 (*Validity:* invalid, valid cue) × 2 (*Conflict:* congruent, incongruent) was conducted to examine the validity × conflict interaction separately in each hemisphere. Effects sizes are reported as partial eta squared (h^2^).

## Results

**Table 2** shows the RT and error rate for all conditions, separately for each hemisphere. The overall mean RT was 598 ms (SD = 95 ms), and the overall mean error rate was 9.12% (SD = 9.90). Participants were significantly faster at responding to stimuli presented to the RH (598 ± 93 ms) compared to the LH (602 ± 96 ms, *t*(47) = −2.71, *p* < .05, *d* = 0.04), while this difference was not significant for the error rate (RH: 9.02 ± 10.00%, LH: 9.22 ± 9.82%, *t*_(47)_ = −0.42, *p* = .97, *d* = 0.02).

**Table 2.** Mean reaction time in ms (± SD) and Error Rate in percentage (± SD) for each task condition and separately for the two hemispheres

| RT | Congruent |  |  |  | Incongruent |  |  |  |
| --- | --- | --- | --- | --- | --- | --- | --- | --- |
|  | nocue | double | valid | invalid | nocue | double | valid | invalid |
| R | 563 | 529 | 504 | 568 | 687 | 634 | 590 (90) | 719 |
| H | (67) | (70) | (66) | (73) | (112) | (111) |  | (118) |
| L | 570 | 531 | 493 | 573 | 702 | 675 | 606 (96) | 720 |
| H | (87) | (68) | (61) | (78) | (109) | (107) |  | (108) |
| ER |  |  |  |  |  |  |  |  |
| R | 2.95 | 1.56 | 2.43 | 3.82 | 14.93 | 11.28 | 10.13 | 23.09 |
| H | (5.83) | (3.71) | (4.27) | (6.19) | (14.17) | (12.45) | (10.53) | (18.05) |
| L | 4.17 | 2.95 | 1.80 | 2.26 | 15.80 | 14.41 | 9.20 | 19.79 |
| H | (6.66) | (4.70) | (2.40) | (4.46) | (13.12) | (15.54) | (9.54) | (17.58) |

### Hemispherical differences in the attentional effects

**Figure 2** shows the alerting, orienting, disengaging, validity, and conflict effects separately for each hemisphere. For the alerting effect, the hemispherical difference was not significant in RT (RH: 44 ± 37 ms, LH: 33 ± 36 ms, *t*(47) = 1.42, *p* = .08, *d* = .29), as well as in error rate (RH: 2.52 ± 7.23%, LH: 1.30 ± 6.50%, *t*(47) = 1.19, *p* = .12, *d* = .24). For the orienting effect, the hemispherical difference was significant in RT (*t*(47) = 2.66, *p* < .01, *d* = .54), indicating that the orienting effect was smaller when stimuli were presented to the RH (34 ± 28 ms) compared to the LH (53 ± 33 ms). This effect was also significant in error rate (*t*(47) = 2.92, *p* < .001, *d* = .59), with a smaller effect when stimuli were presented to the RH (0.14 ± 5.89%) compared to the LH (3.18 ± 5.73%). For the disengaging, the hemispherical difference was significant in RT (*t*(47) = 2.66, *p* < .01, *d* = .54), indicating the disengaging effect was greater when stimuli were presented to the RH (62 ± 40 ms) compared to the LH (44 ± 34 ms). This effect was also significant in error rate (*t*(47) = 2.93, *p* < .001, *d* = .60), with a greater effect when stimuli were presented to the RH (7.03 ± 8.02%) compared to the LH (2.34 ± 7.03%). For the validity effect, the hemispherical difference was not significant in RT (RH: 96 ± 36 ms, LH: 99 ± 38 ms, *t*(47) = 0.07, *p* = .94, *d* = .01) as well as in error rate (RH: 7.17 ± 7.81%, LH: 5.52 ± 6.71%, (*t*(47) = 1.27, *p* = .21). For the conflict effect (see also **Figure 3**), the hemispherical difference was significant in RT (*t*(47) = .15, *p* < .01, *d* = .03), showing a smaller conflict effect in the RH (106 ± 58 ms) compared to the LH (127 ± 56ms), while this effect was not significant in error rate (RH: 12.17 ± 10.17%, LH: 12.01 ± 11.48%, *t*(47) = 0.14, *p* = .89, *d* = .03).

**Figure 2.**
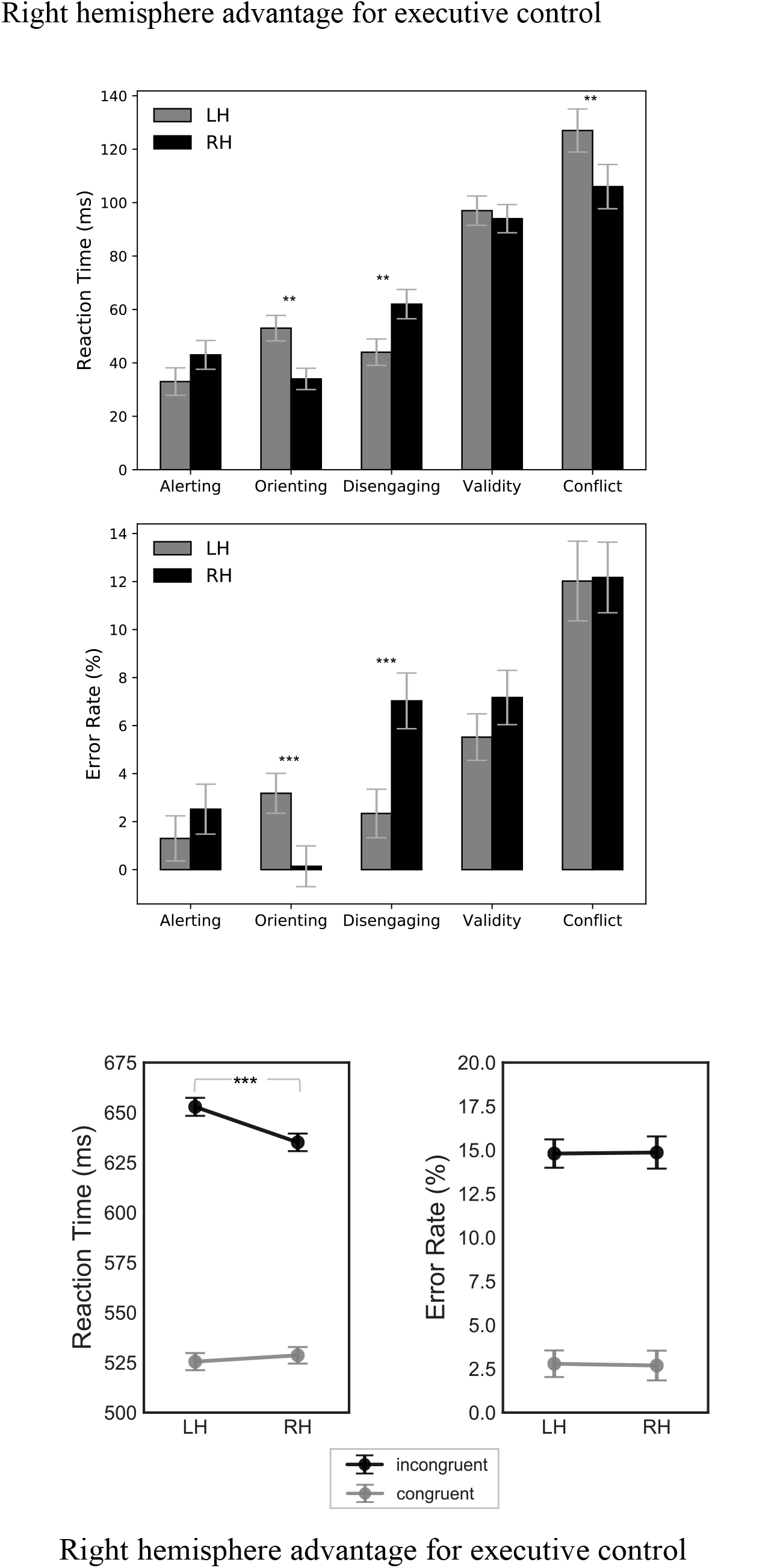
Attentional effects and interactions in terms of reaction time (top panel), and error rate (lower panel). Error bars represent the standard error of the mean. ** = < .01; *** = *p* <.001.

**Figure 3.**
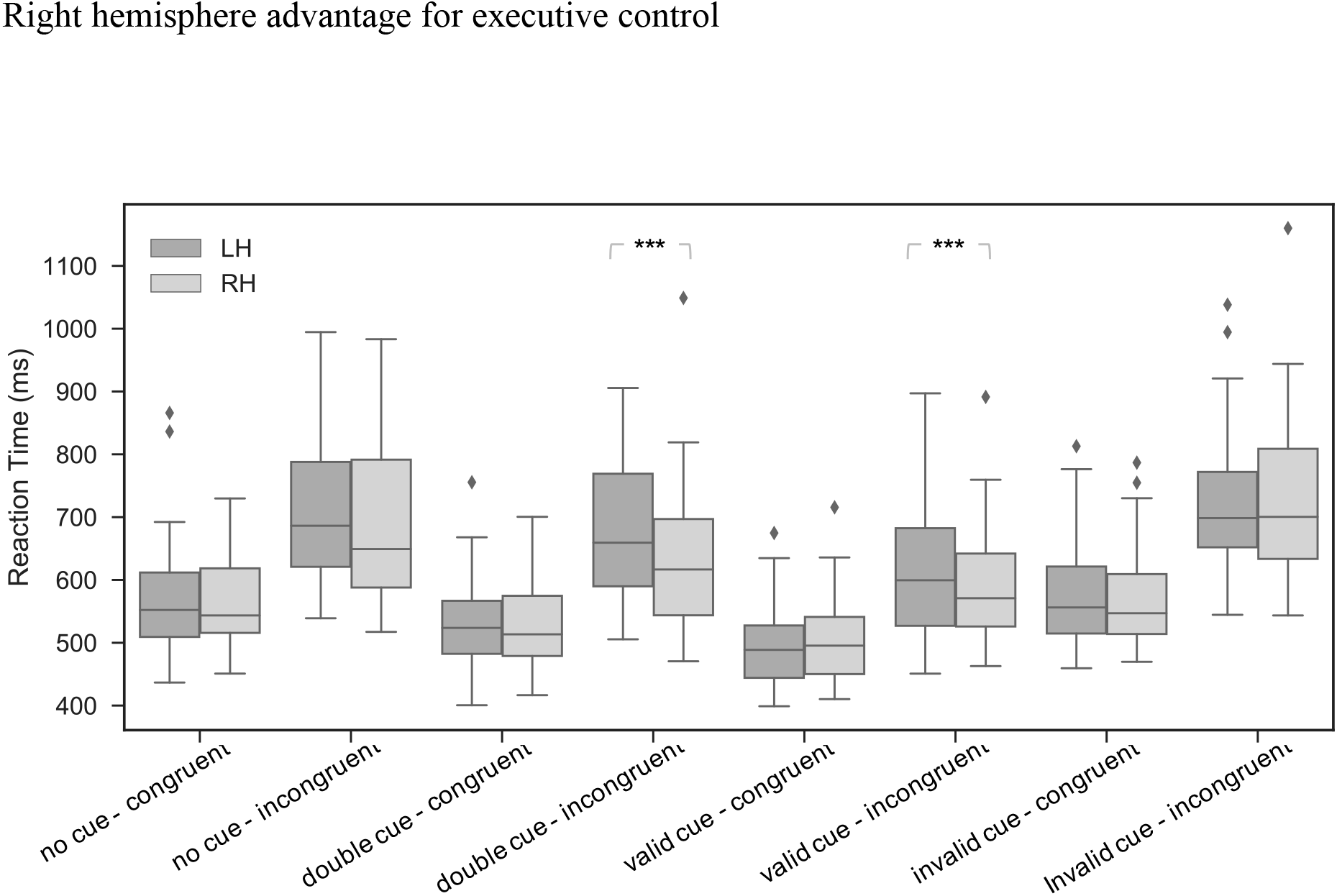
Average reaction time (left panel) and error rate (right panel) for the congruent and incongruent conditions. Error bars represent the standard error of the mean. *** = *p* *.001.

### Hemispherical differences in terms of main effects and interactions

For the analysis conducted in RT, (the top panel of **Figure 4**), the main effect of the factor *Cue* was significant, (*F*(1, 47) = 233.61, *p* < .001, h^2^ = .83), indicating that participants were faster in the valid cue condition (548 ± 94 ms) compared to the double cue (592 ± 111 ms, *p* < .001), nocue (631 ± 114 ms, *p* < .001), and invalid cue (645 ± 121 ms, *p* < .001). Further, participants were faster in the double cue condition compared to the no cue condition (*p* < .001) and invalid cue condition (*p* < .001), and faster in the nocue compared to the invalid cue condition (*p* < .05). The main effect of the factor *Conflict* was significant (*F*(1, 47) = 218.89, *p* < .001, h^2^ = .82), indicating that participants were faster in the congruent condition (541 ± 77 ms) compared to the incongruent condition (667 ± 116 ms, *p* < .001). The main effect of the factor *Hemisphere* was significant (*F*(1, 47) = 7.32, *p* < .01, h^2^ = .14), indicating that participants were faster in response to stimuli presented to the RH (599 ± 114 ms) compared to stimuli presented to the LH (609 ± 119 ms, *p* < .01). The interaction *Cue by Conflict* was significant, (*F*(3, 141) = 15.10, *p* < .001, h^2^ = .24), the interaction *Cue by Hemisphere* was significant (*F*(3, 141) = 3.64, *p* < .05, h^2^ = .07), the interaction *Conflict by Hemisphere* was significant (*F*(1, 47) = 13.14, *p* < .001, h^2^ = .22), and the three way interaction *Cue by Conflict by Hemisphere* was also significant (*F*(3, 141) = 4.26, *p* < .01, h^2^ = .08). In line with the hypothesis driven nature of the current manuscript, planned comparisons were conducted to further analyze these interactions (see section below).

**Figure 4.**
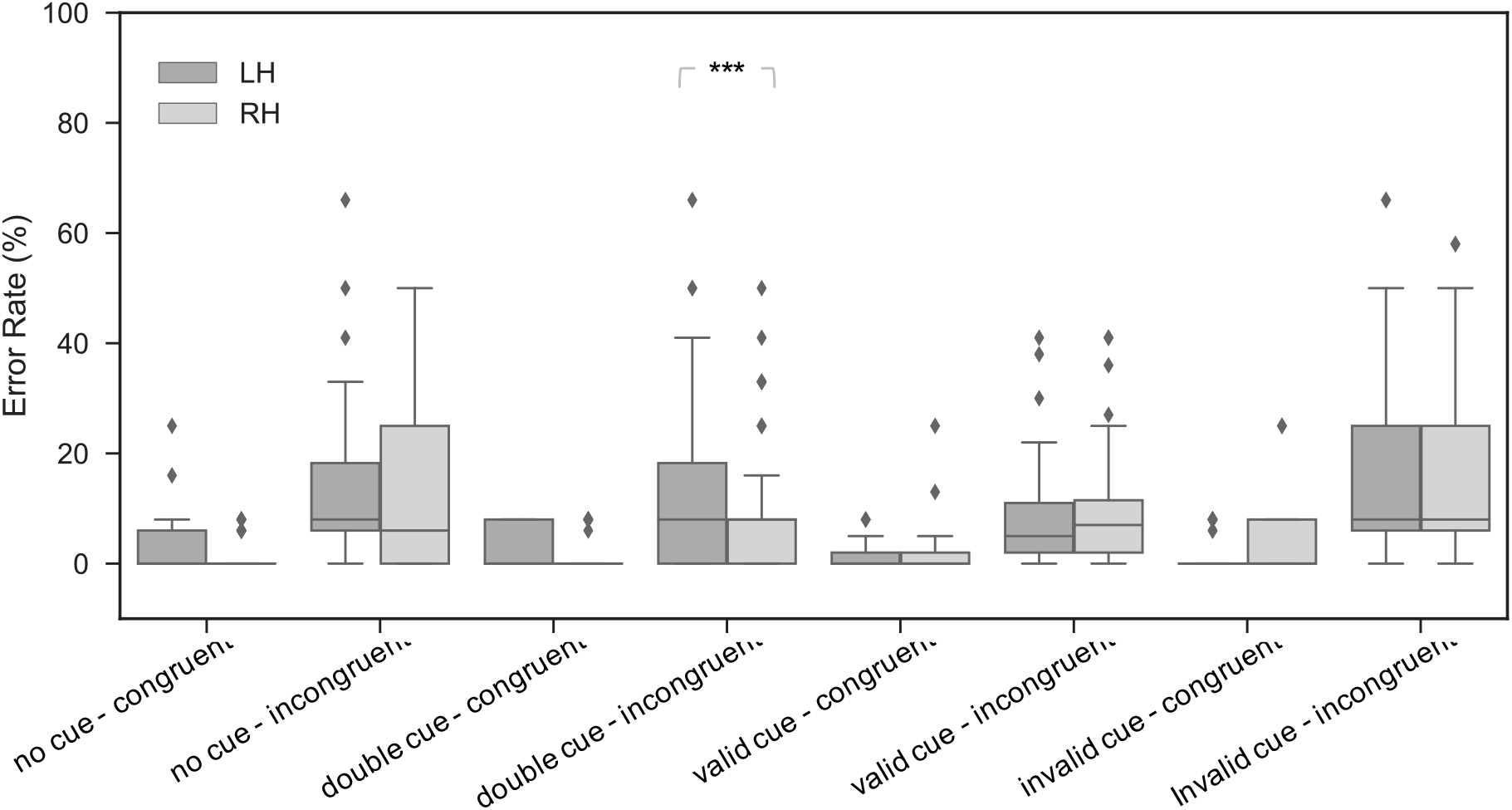
Average reaction time (top panel) and error rate (lower panel) for the four cue conditions: a) no cue; b) double cue; c) valid cue; d) invalid cue, separately for the incongruent and congruent conditions and for stimuli presented in the RH or LH. Error bars representstandard error of the mean.

For the analysis conducted in Error rate, (the lower panel of **Figure 4**), the main effect of the factor *Cue* was significant, (*F*(1, 47) = 24.95, *p* < .001, h^2^ = .35), indicating that participants committed fewer errors in the valid cue condition (5.89 ± 8.37%) compared to the nocue (9.46 ± 12.10%, *p* < .001), and invalid cue (12.24 ± 16.04 ms, *p* < .001), while the difference was not significant compared to the double cue (7.55 ± 11.67%, *p* = .09). Further, participants made fewer errors in the double cue condition compared to the invalid cue condition (*p* < .001) while this difference was not significant compared to the no cue condition (*p* = .17). The difference between nocue and invalid cue trials was significant (*p* < .05). The main effect of the factor *Conflict* was significant (*F*(1, 47) = 68.58, *p* < .001, h^2^ = .59), indicating that participants made fewer errors in the congruent condition (2.74 ± 4.98%) compared to the incongruent condition (14.83 ± 14.74%, *p* < .001). The main effect of the factor *Hemisphere* was not significant (*F* <1). The interaction *Cue by Conflict* (*F*(3, 141) = 21.09, *p* < .001, h^2^ = .31) and the *Cue by Hemisphere* interactions (*F*(3, 141) = 5.05, *p* < .01, h^2^ = .10) were significant, while the *Conflict by Hemisphere* (*F*<1) and the three way interaction *Cue by Conflict by Hemisphere* (*F* <1) were not. In line with the hypothesis driven nature of the current manuscript, planned comparisons were conducted to further analyze these interactions (see section below).

### Hemispherical differences in terms of the interactions between attentional functions

For brevity, here we report only results of the three-way interactions between the conflict effect and the hemispheres under different cue conditions, which are the focus of the current paper. A full depiction of the results regarding the 2 (*Hemisphere*) x 2 (*Cue conditions*) x 2 (*Conflict*) ANOVAs can be found in **Figure 5**, and a full description can be found in **Table S1**.

**Figure 5.**
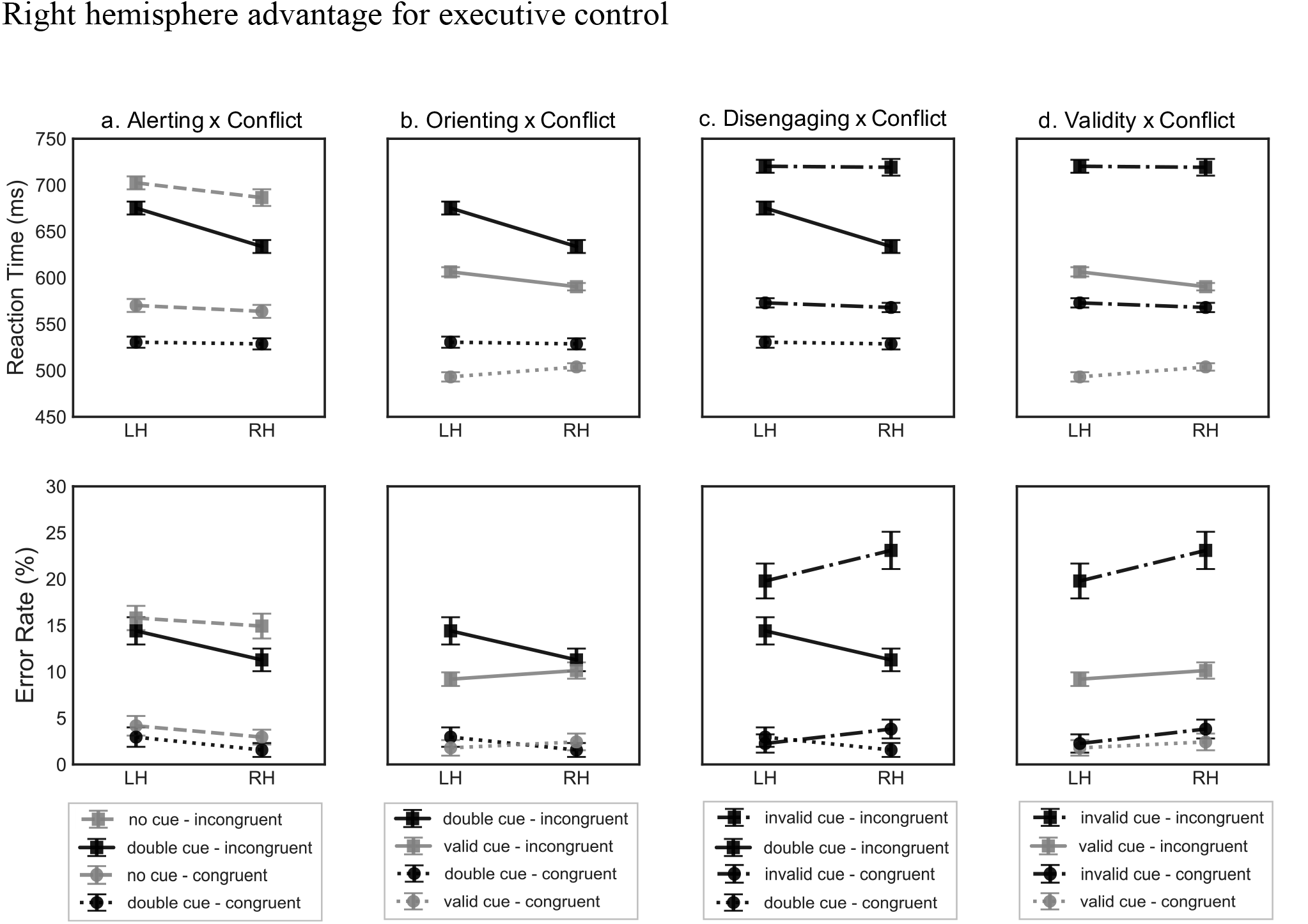
Average reaction time (top panel) and error rate (lower panel) for the interactions between the task conditions: a) alerting by conflict; b) orienting by conflict; c) disengaging by conflict; d) validity by conflict. Error bars represent the standard error of the mean.

*The Alerting by Conflict interaction.* For the 2 (*Hemisphere:* RH, LH) x 2 (*Alerting:* no cue, double cue) x 2 ANOVA (*Conflict:* congruent, incongruent) conducted on the RT (the top panel of **Figure 5a**), the three-way interaction was significant (*F*(1, 47) = 4.03, *p* = .05, h^2^ = .08). The follow-up analysis conducted on the *Hemisphere* by *Conflict* interaction separately for the no cue and double cue conditions showed that for the no cue condition, showed that the twoway interaction was not significant (*F* < 1), while for the double cue condition the two-way interaction was significant (*F*(1, 47) = 18.49, *p* < .001, h^2^ = .28), showing that participants were significantly faster at the incongruent condition when stimuli were presented to the RH (634 ± 111 ms) compared to the LH (675 ± 107 ms, *p* < .001). For the ANOVA conducted on error rate (the bottom panel of **Figure 5a**), the three-way interaction was not significant (*p* ≥ .24).

*The Orienting by Conflict interaction.* For the 2 (*Hemisphere:* RH, LH) x 2 (*Alerting:* double cue, valid cue) x 2 ANOVA (*Conflict:* congruent, incongruent) conducted on the RT (the top panel of **Figure 5b**), the three-way interaction was not significant (*F*(1, 47) = 1.33, *p* = .25, h^2^ = .03). For the ANOVA on error rate (the bottom panel of **Figure 5b**), the three-way interaction was also not significant (*p* ≥ .38).

*The Disengaging by Conflict interaction.* For the 2 (*Hemisphere:* RH, LH) × 2 (*Alerting:* double cue, invalid cue) × 2 ANOVA (*Conflict:* congruent, incongruent) conducted on the RT (the top panel of **Figure 5c**), the three-way interaction was significant (*F*(1, 47) = 15.38, *p* < .001, h^2^ = .25). The follow-up analysis conducted on the *Hemisphere* by *Conflict* interaction separately for the invalid cue and double cue conditions showed that for the invalid cue condition, the two-way interaction was not significant (*F* < 1), while it was significant for the double cue condition, (*F*(1, 47) = 18.49, *p* < .001, h^2^ = .28), showing that participants were significantly faster at the incongruent condition when stimuli were presented to the RH (634 ± 111 ms) compared to the LH (675 ± 107 ms, *p* < .001), while this difference was not significant for the congruent condition (RH: 529 ± 70 ms, LH: 531 ± 68ms, *p* = .74). For the ANOVA conducted on error rate (the bottom panel of **Figure 5c**), the three-way interaction was not significant (*p* ≥ .36).

*The Validity by Conflict interaction.* For the 2 (*Hemisphere:* RH, LH) x 2 (*Alerting:* double cue, invalid cue) x 2 ANOVA (*Conflict:* congruent, incongruent) conducted on the RT (the top panel of **Figure 5d)**, the three-way interaction was significant (*F*(1, 47) = 7.27, *p* < .01, h^2^ = .13). The follow-up analysis conducted on the *Hemisphere* by *Conflict* interaction separately for the valid cue and invalid cue conditions showed that for the valid cue condition, the two-way interaction was significant (*F*(1, 47) = 25.23, *p* < .001, h^2^ = .35), indicating that for the congruent condition participants were slower in the RH (503 ± 66 ms) compared to the LH (493 ± 61 ms), while in the incongruent condition participants were faster in the RH (590 ± 90 ms) compared to the LH (606 ± 95 ms). For the invalid cue condition, the two-way interaction was not significant (*F* < 1). For the ANOVA conducted on error rate (the bottom panel of **Figure 5c**), the three-way interaction was not significant (*p* ≥ .36).

## Discussion

In the present study, we used a lateralized version of the ANT-R and the visual field methodology to examine the efficiency and interactions of the attentional networks separately in the RH and LH, with a specific focus to the hemispherical asymmetry of the executive control under different alerting and orienting conditions. In summary, significant behavioral differences were found for the orienting, disengaging, and executive control of attention, and interactions between either the alerting and the orienting function with the executive control of attention, showing an advantage in all these functions for the RH. These result supports the long-standing proposal of a RH advantage for the orienting of attention and extend it to also include hemispherical asymmetries for executive control of attention. Specifically, the RH advantage in conflict resolution occurs under conditions of increased alertness or valid spatial orienting, suggesting a synergistic dynamic among the three functions. Nonetheless, our results showed no evidence for a hemispherical difference in the spatial cue conditions (i.e., no difference between left or right cues for either valid or invalid conditions). Therefore, we have no evidence for a “disadvantage” of the LH compared to the RH in the orienting of attention (a result that would have favored the hemispatial theory (Heilman & Van Den Abell, 1980), and while it is plausible that, in line with the interhemisperic competition account (Kinsbourne, 1987), each hemisphere has its own contralateral vector of attention, although the lack of lesion data makes our conclusion speculative.

Consistent with our (Fan et al., 2009; Mackie et al., 2013; Spagna, Mackie, et al., 2015) and other studies (Asanowicz & Marzecová, 2017; Callejas et al., 2005; Callejas et al., 2004; Chica et al., 2012; Marotta et al., 2012; Roca et al., 2012), a complex pattern of interactions was shown among the three attentional functions, further proving that the interplay among them is necessary for the selection and prioritization of the processing of goal-relevant information (Badre, 2011). For instance, we found that indicating the location where the target would be presented by means of a valid spatial cue reduced the conflict effect by decreasing the response time to a target flanked by incongruent information. This is consistent with previous evidence showing the beneficial effect of the endogenous orienting function on the executive control of attention (e.g., Callejas et al., 2005; Fan et al., 2009; Spagna, Mackie, et al., 2015; Xuan et al., 2016), and our results further extend this knowledge to the exogenous orienting function, by showing that the conflict resolution benefited more from a valid cue compared to an invalid cue when the process was conducted by the RH. Conversely, a wealth of studies showed that the presentation of an alerting cue has a cost on the conflict resolution (Asanowicz & Marzecová, 2017; Callejas et al., 2005; Callejas et al., 2004; Xuan et al., 2016), a pattern theoretically linked to a “U-shape” relationship (and justified by the norepinephrine dose-response relationship underlying the alerting function (Aston-Jones & Cohen, 2005)), with a phasic increase of the arousal state altering the efficiency of the executive control of attention (Thiele & Bellgrove, 2018). In this study, however, we show a pattern of facilitation produced by an alerting cue on the conflict resolution when stimuli were processed by the RH compared to the LH, indicating the superiority of this hemisphere in conflict resolution is benefited by the activation of the alerting function. At a closer look, our results and the evidence from the previous studies can coexist due to the following reasons: 1) we also show a greater conflict effect in the double cue condition compared to the no cue condition, when averaging performance across the two hemispheres; 2) however, if further broken down into the specific task conditions, the above mentioned effect reveals that participants showed overall better performance at the double cue compared to the no cue effect; and 3) the interaction between these two components of attention within each visual field shows that a double cue condition was significantly less beneficial to the conflict processing in the LH. While the most obvious explanation for this interaction effect is to state a RH advantage for both alerting, executive control, and their interaction, it may also be that the worsening of the conflict resolution in the LH under the alerting condition results from more limited attentional resources of this hemisphere that more quickly are exhausted when the two functions are contextually engaged.

Consensus exists regarding the neural substrate supporting the activation of interactive attentional networks (Xuan et al., 2016), showed as a complex pattern of interaction among cortical and subcortical areas (Petersen & Posner, 2012). Specifically, a bilateral fronto-parietal network (composed of the frontal eye fields, areas near and along the intraparietal sulcus, and the dorsolateral prefrontal cortex), a cingulo-opercular network (composed of the anterior cingulate cortex, and the anterior insular cortex), and subcortical areas such as the thalamus, superior colliculi, and the basal ganglia, were found to consistently interact during attentional tasks (Callejas, Shulman, & Corbetta, 2014; Corbetta et al., 2008; Corbetta & Shulman, 2002; Fan et al., 2005; Krauzlis, Lovejoy, & Zénon, 2013; Parlatini et al., 2017; Patel et al., 2015; Posner, 2012; Posner & Fan, 2008; Shulman et al., 2009; Wang et al., 2010; Xuan et al., 2016). However, bilateral involvement and functional asymmetries are not mutually exclusive properties in our brain (Arbula et al., 2017; Bartolomeo & Thiebaut de Schotten, 2016; Corballis, 2009, 2017; Vallesi, Arbula, Capizzi, Causin, & D’Avella, 2015), and the existence of hemispherical differences for executive control of attention in terms of activation magnitude seems plausible. This is particularly true if we consider that the function of executive control of attention is to allow the selection and prioritization of the processing of goal-relevant information to reach consciousness (Fan, Fossella, Sommer, Wu, & Posner, 2003; Mackie & Fan, 2017; Mackie et al., 2013; Petersen & Posner, 2012; Spagna, Mackie, et al., 2015) and that this function inevitably interacts with the hemispherical specializations related to the specific type of information to be handled (Marotta & Casagrande, 2017; Spagna et al., 2016; Spagna et al., 2014). More specifically, hemispherical asymmetries were first identified in “split brain” patients (i.e., corpus calloscotomy intervention as part of medical treatment) (Sperry, 1968), one of the earliest evidence for hemispheric dominance for verbal and language functions in the LH and for non-verbal and spatial functions in the RH (see Bartolomeo & Thiebaut de Schotten, 2016 for a review). In the present study, we used non-verbal information as imperative stimuli, an aspect that might have magnified the subtle interactions among the executive control of attention and the hemispheric dominance of the RH for this type of stimuli.

Alternatively, the RH advantage in conflict resolution may be explained by brain mechanisms revealed in neuroimaging studies. Neuroimaging evidence for a RH dominance associated with the executive control of attention exist at the functional level (Garavan et al., 1999), as shown by the association between participants’ monitoring of task contingencies and the activation of prefrontal cortex in the RH (see Vallesi, 2012 for a review; Vallesi & Crescentini, 2011), response inhibition and the association with the right inferior frontal cortex (Aron, Robbins, & Poldrack, 2014; Cai, Ryali, Chen, Li, & Menon, 2014; Levy & Wagner, 2011), increased metabolic activity in the medial prefrontal cortex associated with monitor the auditory environment (Cohen, Semple, Gross, King, & Nordahl, 1992), and the anterior cingulate cortex of the RH for response selection (Turken & Swick, 1999), motor control (Paus, 2001) and attention shifting (Kondo, Osaka, & Osaka, 2004). Further, regions within the frontal and parietal lobes of the right hemisphere have been consistently shown to play a critical role in the more broad construct of attention, by modulating activity in the sensory cortices (Bressler, Tang, Sylvester, Shulman, & Corbetta, 2008; Ruff et al., 2009), and by dynamically adjusting the focus of attention (Malhotra, Coulthard, & Husain, 2009; Marshall, O’Shea, Jensen, & Bergmann, 2015; O’Shea, Muggleton, Cowey, & Walsh, 2004; Ronconi, Basso, Gori, & Facoetti, 2014). Consistently, in one of our previous studies using the ANT together with fMRI (Fan et al., 2005), in which the presentation of the stimuli was not lateralized, we showed a significant activation of the right temporo-parietal junction for the alerting function, together with clusters in the right parietal lobe for the orienting function, and a right-centered cluster of anterior cingulate cortex activation for executive control of attention. Together with the evidence regarding the necessity of the fronto-parietal and cingulo-opercular networks of the RH for both the alerting (J. Li et al., 2016; Q. Li et al., 2018; Perin, Godefroy, Fall, & de Marco, 2010; Posner, 2008) and orienting (Bartolomeo, 2014; Bartolomeo, Thiebaut de Schotten, & Chica, 2012; Chica et al., 2018; Chica et al., 2012; Mesulam, 1999), this neuroimaging evidence is in support of the findings from the present study regarding the executive control of attention.

One limitation of the present study must be, however, kept in mind. Here, we did not employ eye tracking techniques and therefore have no direct measure of the eye position of participants throughout the task. This is point may limit the conclusion of our studies, because eye movements must be avoided at all costs for the lateralized presentation to work. However, evidence exists that suggests carefully instructing participants before the beginning of the task regarding the relevance of maintaining central fixation can be effective in controlling eye movements (e.g., Moscovitch, 1986; Śmigasiewicz et al., 2014; Smigasiewicz et al., 2010; Verleger, Smigasiewicz, & Moller, 2011; Verleger et al., 2009). Additionally, a variety of behavioral studies investigating the hemispherical differences in the attentional functions using behavioral tasks with lateralized stimuli found interactions between the attentional functions and the visual field with and without using eye-tracker. It is worth noting that the lack of a control of eye movement may have a negative impact so that the hemispheric difference would not be significant. Here, we are not making conclusions about the negative findings (absence of hemispherical advantage) but on the positive findings (presence of hemispherical advantage of the RH). In other words, the significant difference between hemisphere should not be due to the presence of eye movement, but due to the true effect in terms of hemispherical difference. Overall, although the lack of control of participants’ central fixation limits the validity of our conclusion regarding the orienting effect, it is not as relevant for our result concerning the executive control of attention, which was the focus of the study.

Our study is consistent with existing behavioral and neural evidence indicating a RH advantage for attention, and further provides novel knowledge regarding how this advantage also extends to the executive control of attention and its dynamics with the alerting and orienting function. These results may shed light on some of the inconsistencies shown in previous studies that did not find RH dominance for executive control and for the interplay between the three functions and pave the way for future studies by combining our task with neuroimaging techniques, aimed at characterizing the neural dynamics underlying the RH advantage for executive control of attention.

## Acknowledgements

This study was supported by the National Institute of Mental Health (NIMH) of the National Institutes of Health (NIH) under Award Number R01 MH094305.

---

**Table S1.**
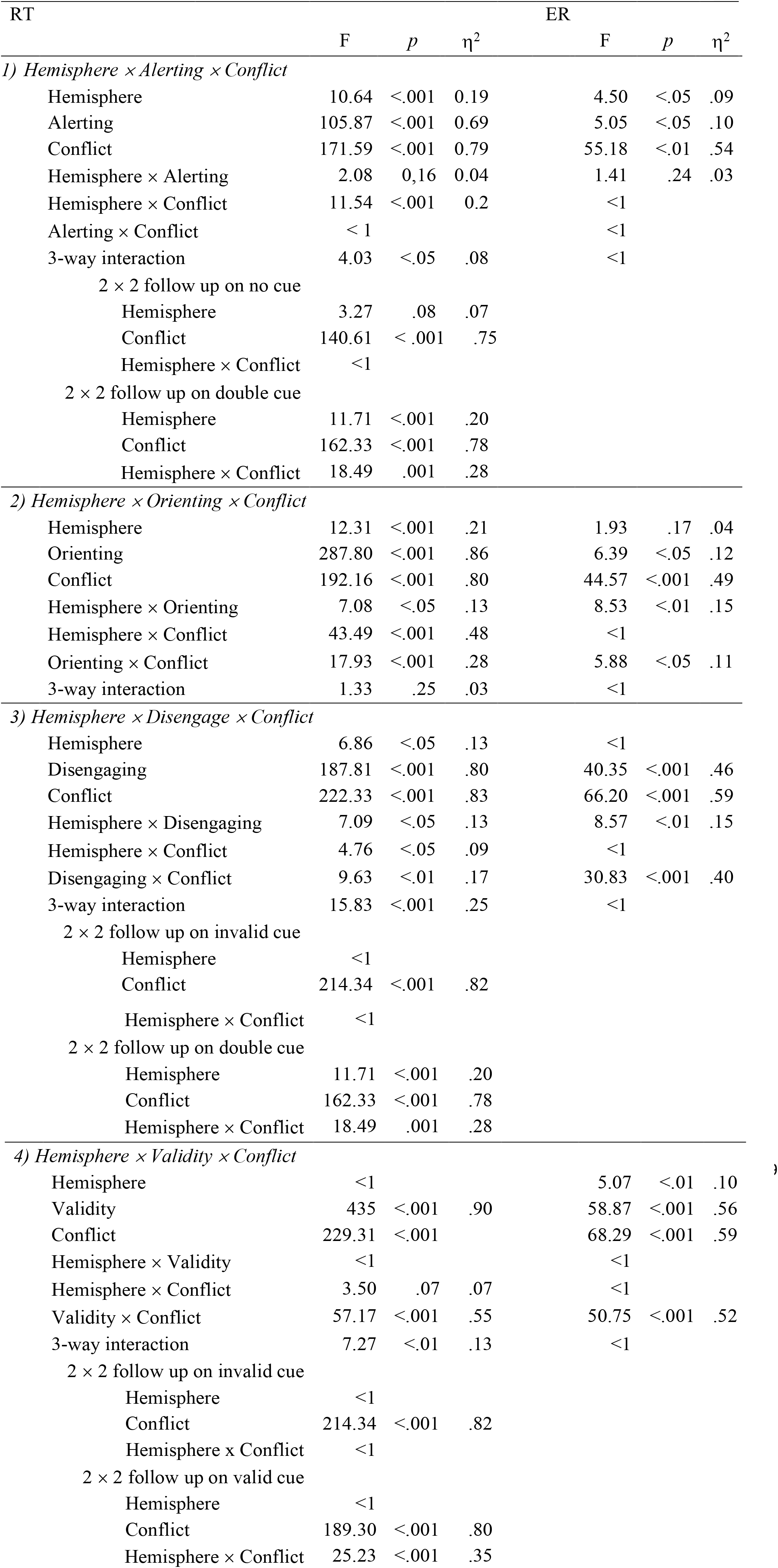
Results of the ANOVAs on the hemispherical differences in terms of the interactions between attentional functions

| RT |  |  |  | ER |  |  |
| --- | --- | --- | --- | --- | --- | --- |
| | F | *p* | $\eta^2$ | F | *p* | $\eta^2$ |
| *1) Hemisphere × Alerting × Conflict* |  |  |  |  |  |  |
| Hemisphere | 10.64 | <.001 | 0.19 | 4.50 | <.05 | .09 |
| Alerting | 105.87 | <.001 | 0.69 | 5.05 | <.05 | .10 |
| Conflict | 171.59 | <.001 | 0.79 | 55.18 | <.01 | .54 |
| Hemisphere × Alerting | 2.08 | 0.16 | 0.04 | 1.41 | .24 | .03 |
| Hemisphere × Conflict | 11.54 | <.001 | 0.2 | <1 |  |  |
| Alerting × Conflict | < 1 |  |  | <1 |  |  |
| 3-way interaction | 4.03 | <.05 | .08 | <1 |  |  |
| 2 × 2 follow up on no cue |  |  |  |  |  |  |
| Hemisphere | 3.27 | .08 | .07 |  |  |  |
| Conflict | 140.61 | <.001 | .75 |  |  |  |
| Hemisphere × Conflict | <1 |  |  |  |  |  |
| 2 × 2 follow up on double cue |  |  |  |  |  |  |
| Hemisphere | 11.71 | <.001 | .20 |  |  |  |
| Conflict | 162.33 | <.001 | .78 |  |  |  |
| Hemisphere × Conflict | 18.49 | .001 | .28 |  |  |  |
| *2) Hemisphere × Orienting × Conflict* |  |  |  |  |  |  |
| Hemisphere | 12.31 | <.001 | .21 | 1.93 | .17 | .04 |
| Orienting | 287.80 | <.001 | .86 | 6.39 | <.05 | .12 |
| Conflict | 192.16 | <.001 | .80 | 44.57 | <.001 | .49 |
| Hemisphere × Orienting | 7.08 | <.05 | .13 | 8.53 | <.01 | .15 |
| Hemisphere × Conflict | 43.49 | <.001 | .48 | <1 |  |  |
| Orienting × Conflict | 17.93 | <.001 | .28 | 5.88 | <.05 | .11 |
| 3-way interaction | 1.33 | .25 | .03 | <1 |  |  |
| *3) Hemisphere × Disengage × Conflict* |  |  |  |  |  |  |
| Hemisphere | 6.86 | <.05 | .13 | <1 |  |  |
| Disengaging | 187.81 | <.001 | .80 | 40.35 | <.001 | .46 |
| Conflict | 222.33 | <.001 | .83 | 66.20 | <.001 | .59 |
| Hemisphere × Disengaging | 7.09 | <.05 | .13 | 8.57 | <.01 | .15 |
| Hemisphere × Conflict | 4.76 | <.05 | .09 | <1 |  |  |
| Disengaging × Conflict | 9.63 | <.01 | .17 | 30.83 | <.001 | .40 |
| 3-way interaction | 15.83 | <.001 | .25 | <1 |  |  |
| 2 × 2 follow up on invalid cue |  |  |  |  |  |  |
| Hemisphere | <1 |  |  |  |  |  |
| Conflict | 214.34 | <.001 | .82 |  |  |  |
| Hemisphere × Conflict | <1 |  |  |  |  |  |
| 2 × 2 follow up on double cue |  |  |  |  |  |  |
| Hemisphere | 11.71 | <.001 | .20 |  |  |  |
| Conflict | 162.33 | <.001 | .78 |  |  |  |
| Hemisphere × Conflict | 18.49 | .001 | .28 |  |  |  |
| 4) Hemisphere × Validity × Conflict |  |  |  |  |  |  |
| Hemisphere | <1 |  |  | 5.07 | <.01 | .10 |
| Validity | 435 | <.001 | .90 | 58.87 | <.001 | .56 |
| Conflict | 229.31 | <.001 |  | 68.29 | <.001 | .59 |
| Hemisphere × Validity | <1 |  |  | <1 |  |  |
| Hemisphere × Conflict | 3.50 | .07 | .07 | <1 |  |  |
| Validity × Conflict | 57.17 | <.001 | .55 | 50.75 | <.001 | .52 |
| 3-way interaction | 7.27 | <.01 | .13 | <1 |  |  |
| 2 × 2 follow up on invalid cue |  |  |  |  |  |  |
| Hemisphere | <1 |  |  |  |  |  |
| Conflict | 214.34 | <.001 | .82 |  |  |  |
| Hemisphere × Conflict | <1 |  |  |  |  |  |
| 2 × 2 follow up on valid cue |  |  |  |  |  |  |
| Hemisphere | <1 |  |  |  |  |  |
| Conflict | 189.30 | <.001 | .80 |  |  |  |
| Hemisphere × Conflict | 25.23 | <.001 | .35 |  |  |  |

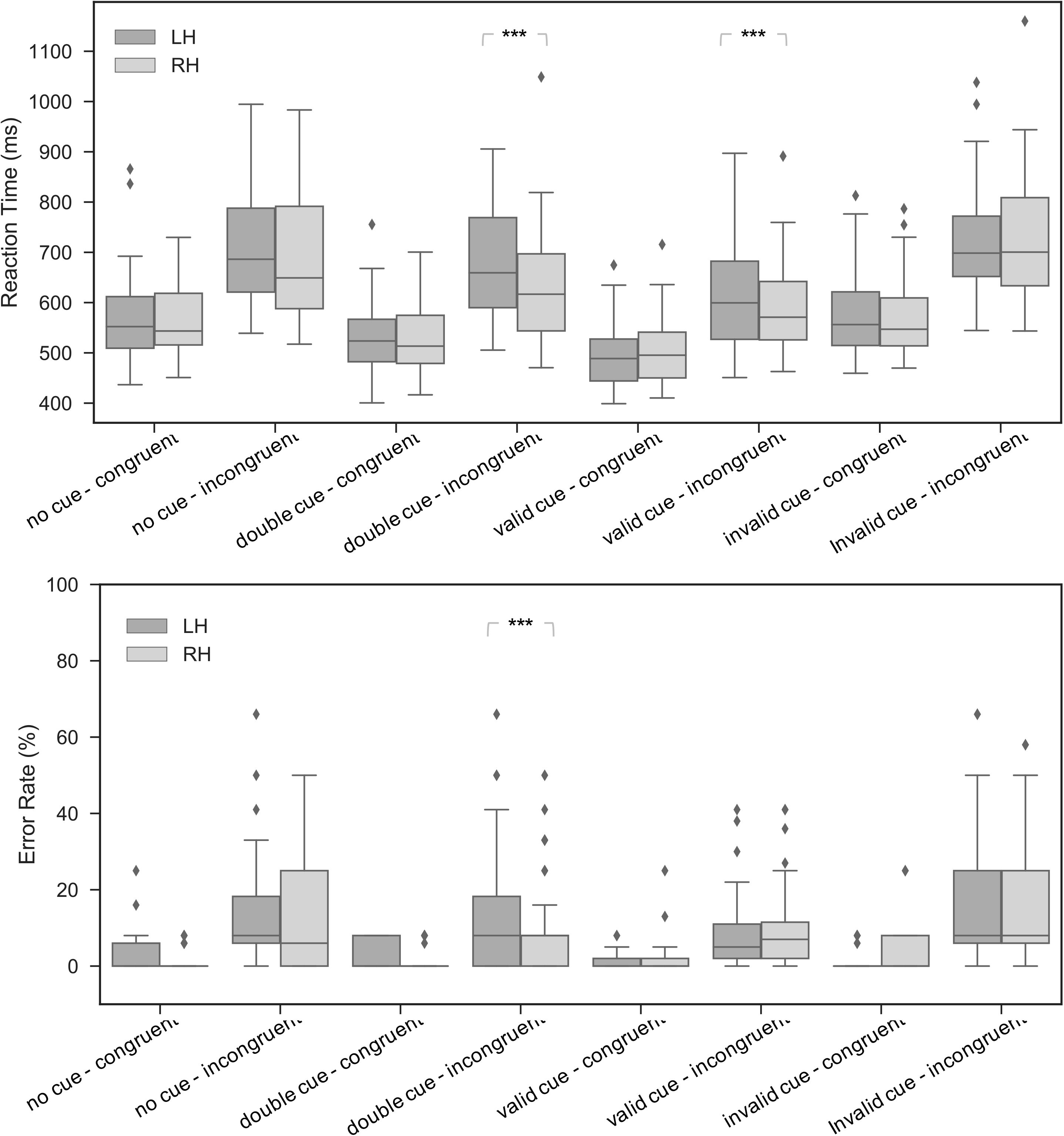

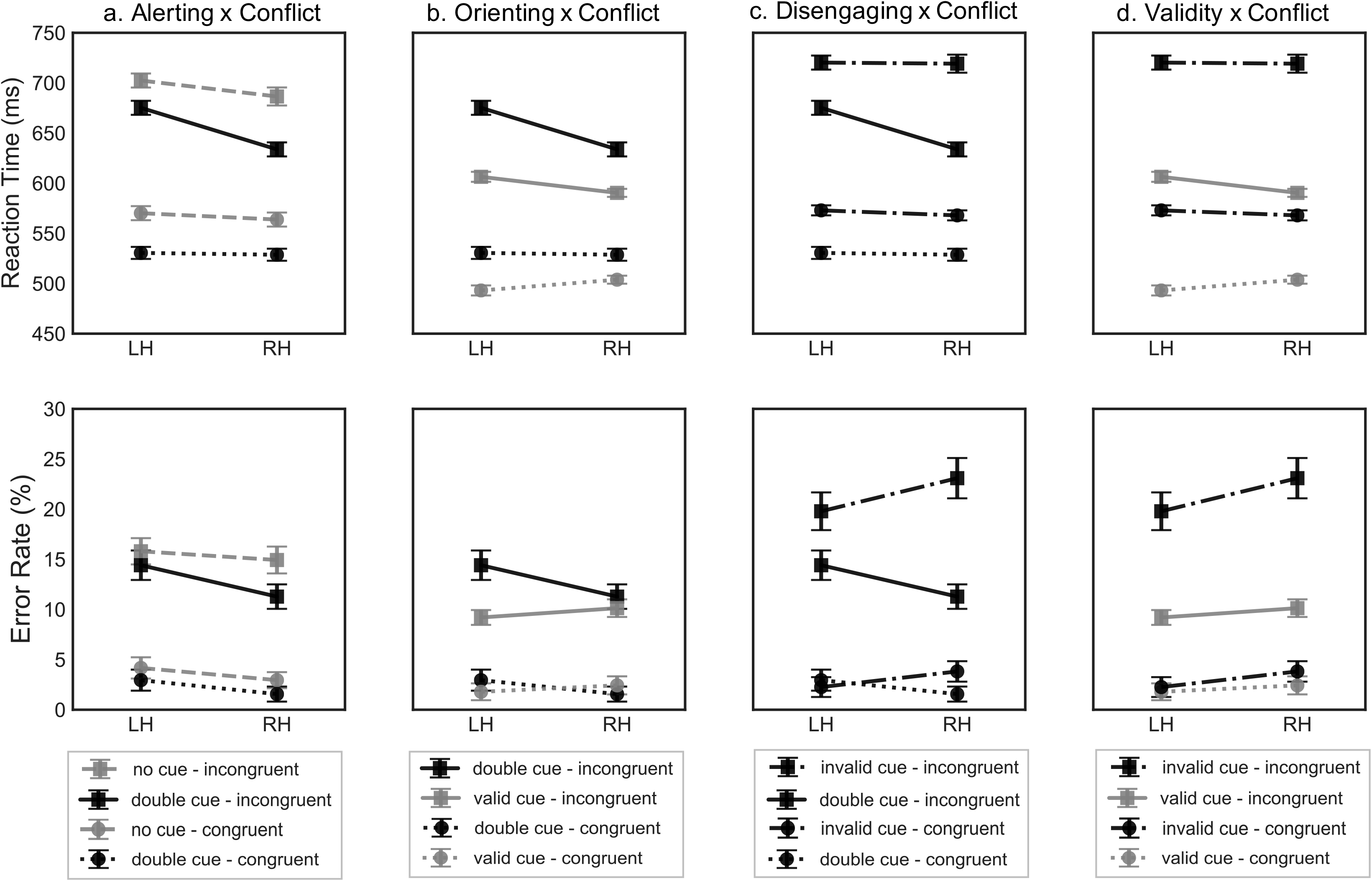

